# Shared neural computations for syntactic and morphological structures: evidence from Mandarin Chinese

**DOI:** 10.1101/2024.01.31.578104

**Authors:** Xinchi Yu, Sebastián Mancha, Xing Tian, Ellen Lau

## Abstract

Although psycho-/neuro-linguistics has assumed a distinction between morphological and syntactic structure building as in traditional theoretical linguistics, this distinction has been increasingly challenged by theoretical linguists in recent years. Opposing a sharp, lexicalist distinction between morphology and syntax, non-lexicalist theories propose common morpho-syntactic structure building operations that cut across the realms of “morphology” and “syntax”, which are considered distinct territories in lexicalist theories. Taking advantage of two pairs of contrasts in Mandarin Chinese with desirable linguistic properties, namely compound vs. simplex nouns (the “morphology” contrast, differing in morphological structure complexity per lexicalist theories) and separable vs. inseparable verbs (the “syntax” contrast, differing in syntactic structure complexity per lexicalist theories), we report one of the first pieces of evidence for shared neural responses for morphological and syntactic structure complexity in language comprehension, supporting a non-lexicalist view where shared neural computations are employed across morpho-syntactic structure building. Specifically, we observed that the two contrasts both modulated neural responses in left anterior and centro-parietal electrodes in an a priori 275:400 ms time window, corroborated by topographical similarity analyses. These results serve as preliminary yet *prima facie* evidence towards shared neural computations across morphological and syntactic structure building in language comprehension.

## 1. Introduction

Both in speech and in text, language comprehension requires inferring and building up the linguistic structures intended by the speaker or writer on the basis of the sensory evidence they provide. According to the traditional lexicalist view, the linguistic structures include for example morphological structures (structures within words with “morpheme” as its unit) and syntactic structures (structures over phrases in a sentence with “word” as their units). Is there really a sharp distinction between “sentence structure” vs. “word structure” (according to lexicalist theories) in language comprehension, or is syntax “all-the-way-down”, removing the need to posit a separate set of word-forming “morphological” operations (according to non-lexicalist theories; Krauska & Lau, 2023)? Although the psycho-/neuro-linguistics literature traditionally studies morphology and syntax separately following the earlier morphology-syntax distinction in theoretical linguistics (Chomsky, 1970; Lapointe, 1980; Williams, 1981), a growing body of literature in theoretical linguistics challenges the idea that so-called “morphological” and “syntactic” rules/operations are qualitatively distinct (Halle & Marantz, 1993; Noyer, 1998; Siddiqi, 2010; Harley, 2014; Embick, 2015; Preminger, 2021). However, there has been little psycho-/neuro-linguistic evidence on this issue (see Krauska & Lau, 2023; Krauska, 2024).

A natural thought then would be to contrast conditions that only differ in morphological structure complexity, and conditions that only differ in syntactic structure complexity, and examine if they elicit similar neural responses. However, there are inevitable and widely known challenges in manipulating linguistic structure without simultaneously altering conceptual structure and/or phonological structure (Pylkkänen, 2019; Călinescu et al., 2023). For example, *red boat* is indeed syntactically more complex than *xkq boat* (where *xkq* is a list of meaningless consonant letters), but the former has a more complex semantic-conceptual structure as well (Bemis & Pylkkänen, 2011, 2013). Among recent work in other languages proposing encouraging solutions to this problem with interesting linguistic properties in those languages (Flick & Pylkkänen, 2020; Law & Pylkkänen, 2021; Matar et al., 2021), in the current paper we take advantage of the desired properties of “separable verbs” in Mandarin Chinese — compound forms whose meanings are non-compositional (e.g. *zao4fan3* means “rebel”, where *zao4* means “manufacture” and *fan3* means “reverse”), but which, unlike compounds in English, can be syntactically separated while preserving their non-compositional meaning. This class of compounds contrasts with another set of “inseparable” verb compounds which are matched in conceptual complexity as well as form complexity but which cannot be separated. At the same time, we also included a second set of conditions in which we compared the response to (inseparable) compound nouns with simplex (i.e. “monomorphemic”) nouns. According to lexicalist theories, separable verbs and inseparable verbs (the syntax contrast) only differs systematically in syntactic structure complexity, while compound nouns and simplex nouns (the morphology contrast) only differs systematically in morphological structure complexity; while non-lexicalist theories would posit that the two types of complexities are qualitatively the same. By comparing the responses across the two pairs of conditions (Figure 1a; Table 1), we can ask whether the neural signature for a “syntactic” contrast is qualitatively similar to a “morphological” contrast in the same participants. Of course, given the limited spatial resolution of EEG (electroencephalography), we cannot guarantee that the neural bases are exactly the same for even the same scalp distributions (which is the case for nearly all neuroimaging methods apart from perhaps single-neuron recording). Here we are only trying to determine if there is *prima facie* evidence for shared neural responses, or if it is the case that even such preliminary evidence is hard to obtain. Our current study should be considered a stepping-stone towards eventually answering this research question instead of the final say.

**Figure 1.**
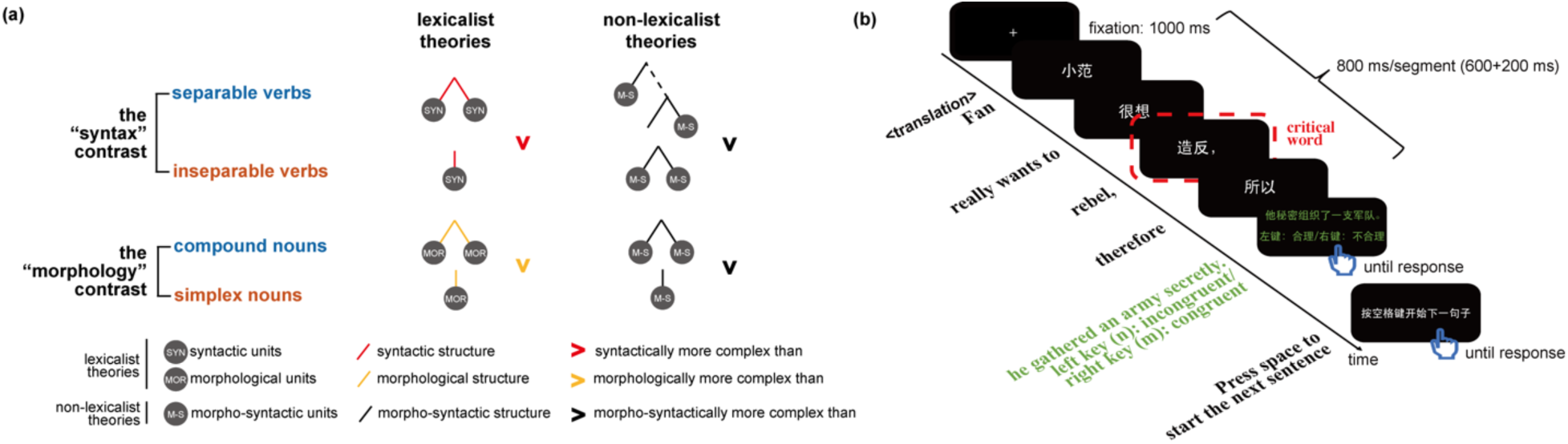
**(a)** Illustration of the two pairs of conditions (i.e., two contrasts) that we employed in the current study. According to lexicalist theories, separable verbs are more complex than inseparable verbs in syntactic structure, while compound verbs are more complex than simplex nouns in morphological structure. Meanwhile, according to non-lexicalist theories, these two complexities are not qualitatively different (i.e., they are both morpho-syntactic in nature). Note that the exact shape of the trees may differ according to different theoretical linguistic theories, yet the complexity difference in the two contrasts holds true for all major relevant theories that we are aware of. **(b)** An illustration of a trial in our current study. English translation is provided below the time arrow.

**Table 1.**
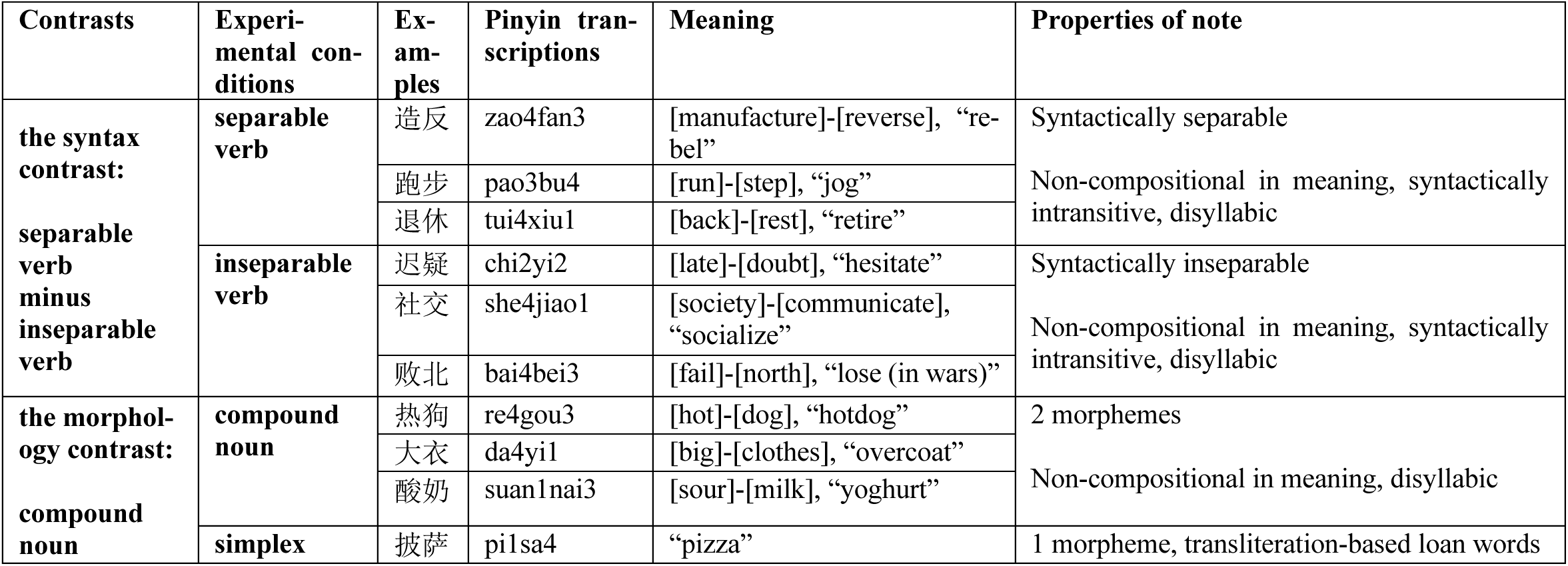
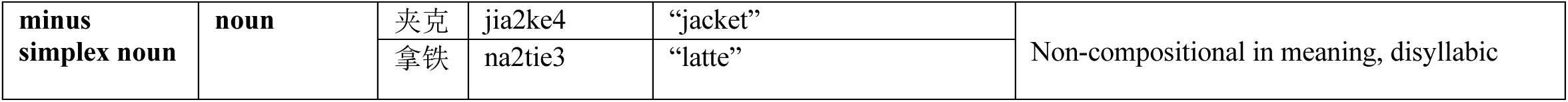
Examples of the separable verbs, inseparable verbs, compound nouns and simplex nouns that we used in the current experiment.

### 1.1. Separable and inseparable verb compounds in Mandarin Chinese

The current study employs an intriguing class of verb compounds in Mandarin Chinese which can be syntactically separated while preserving their non-compositional compound meaning. Here we will use the term “separable verbs”, but in the literature they go by many other names as well, including “ionized forms” (Chao, 1968), “separable words” (Han & Wang, 2022); “splittable compounds” (Siewierska et al., 2010; Lu et al., 2022), as well as its Chinese name *liheci* (离合词, Petrovčič, 2016; Ye & Pan, 2023).

The separable verb phenomenon has been well-documented and extensively discussed in the theoretical linguistics literature (Chao, 1968; Li & Thompson, 1981; Huang, 1984; Packard, 2000; Xue, 2002; Zhang, 2007; Siewierska et al., 2010; Wang, 2011; Chan, 2016; Yeh, 2020; Zhu & Liu, 2020; Ye & Pan, 2023), and somewhat similar phenomena have also been documented in other languages, such as Vietnamese (Nhàn, 1984; Noyer, 1998) and Cantonese (Chan & Cheung, 2021; Yip et al., 2021). An example is illustrated in (1). For the separable verb of “造反” (zao4fan3, [manufacture]-[reverse], “rebel”), syntactic units (e.g., aspect markers, classifiers, quantifiers, resultative verb complements, Siewierska, Xu & Xiao, 2010; Wang, 2011; Ye & Pan, 2023) can be inserted between the two morphemes, but the meaning remains non-compositional (the compound meaning is not transparently derivable as a function of its parts, and thus the compound meaning must be acquired by learners separately from the meaning of its parts). By analogy, if “babysit” in English were a separable compound, one could syntactically separate its parts--e.g., “baby-one time-sit” --while preserving the non-compositional meaning^1^,

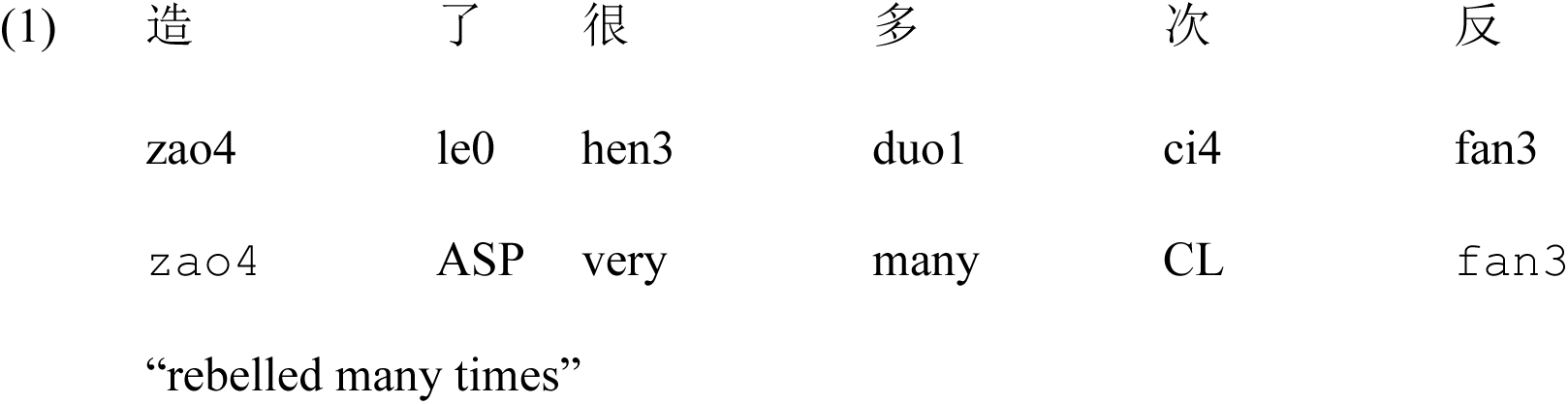

One of the primary reasons that separable verbs have engendered so much interest in the literature is because they seem to put pressure on a traditional notion of “word” as a stored object of semantic, syntactic, and phonological form which enters into semantic, syntactic, and phonological relations with other words. Standard, inseparable compounds like “seahorse” may seem to suggest a non-isomorphic pairing of a single concept to multiple form parts, but these cases can still be accommodated in a traditional framework by assuming that “seahorse” is a single phonological form, which corresponds to a single element in the syntactic structure and which in turn maps to a single concept. Separable verbs cannot be handled this way, because they manifestly demonstrate multiple elements that can be manipulated by the syntax but which realize the single stored (non-compositional) meaning.

The dissociation between conceptual complexity and syntactic complexity evident in Chinese separable verb compounds provides a novel opportunity for investigating the neural correlates of syntactic structure complexity in comprehension, as dissociated from the complexity of conceptual structures. In the current study, we do this by comparing the neural response to separable verb compounds with that of a different syntactic class of verbal compounds which, like English compounds, are “inseparable”. For example, in the case of the inseparable verb of “迟疑” (chi2yi2, [late]-[doubt], “hesitate”), no syntactic units can be inserted in between the two parts, as illustrated in (2a). Instead, one has to use an expression like the one indicated in (2b). For more examples of separable and inseparable verbs we used, see Table 1.

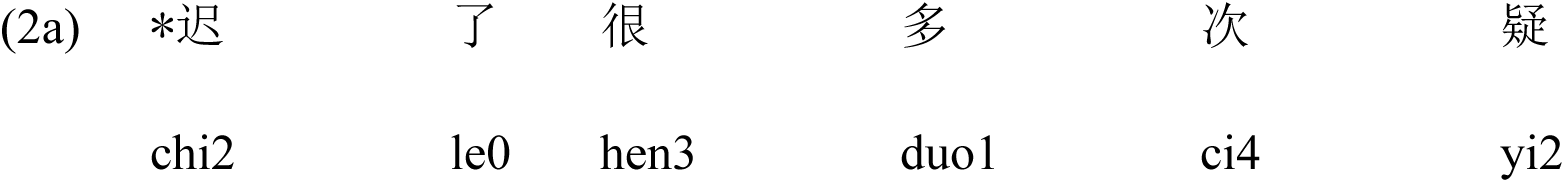
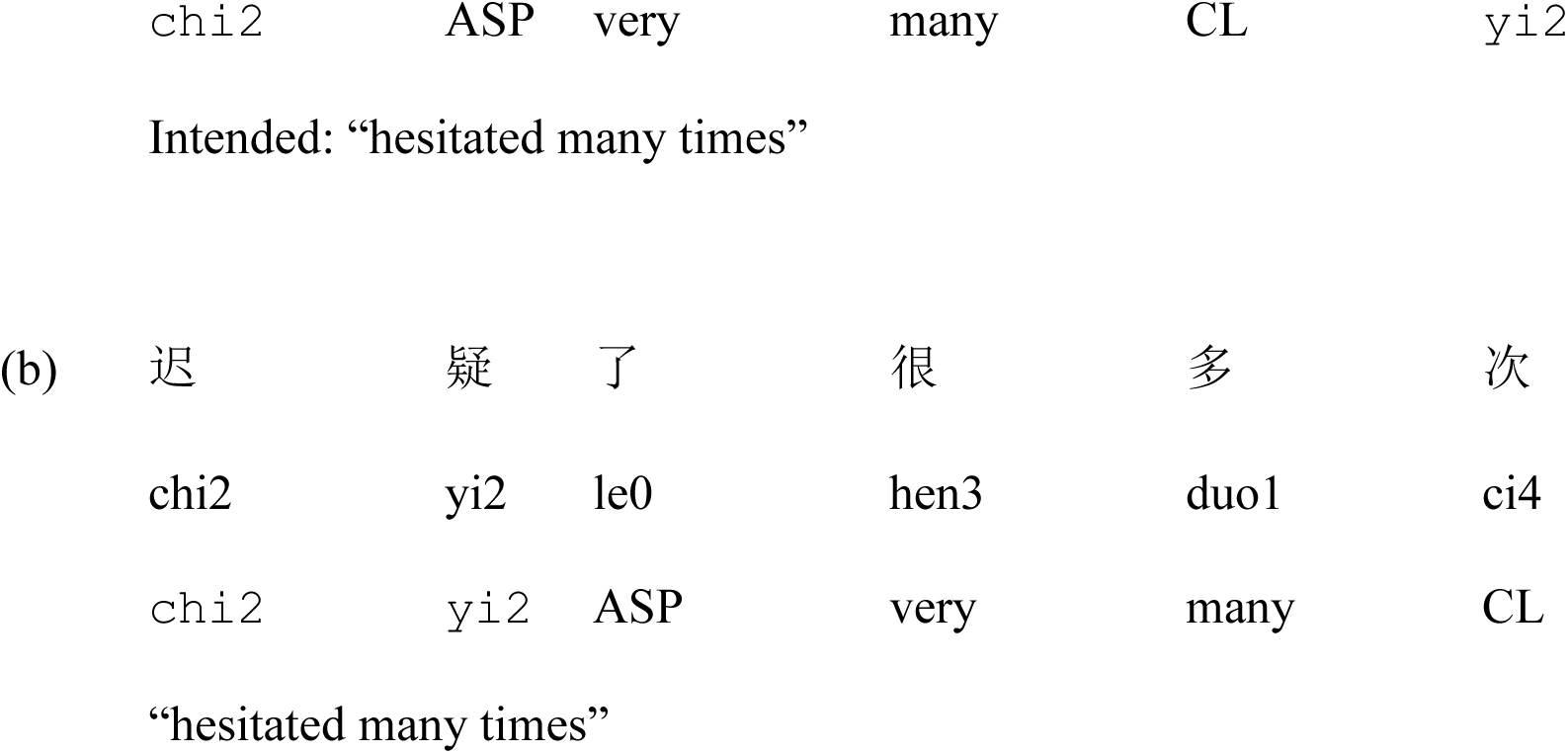

Both separable and inseparable verbs used in the current experiment consist of two morphemes, are non-compositional in meaning, and are syntactically intransitive (i.e., cannot take direct objects syntactically). Therefore, we can assume that the primary systematic difference between the separable vs. inseparable verbs we used here is in their syntactic representation^2^. We note that different syntactic theories vary in exactly how that syntactic difference is realized. In standard lexicalist theories, inseparable compound verbs are considered to consist of a single syntactic unit (i.e., one “word”; Chung, 2006), and separable compound verbs are taken rather to consist of a complex syntactic structure relating multiple syntactic units (Huang, 1997; Badan, 2013; Pan & Ye, 2015; Chan, 2016; Guo, 2017). Non-lexicalist theories, on the other hand, allow the possibility that inseparable compound verbs also have internal syntactic parts corresponding to their compound phonological form parts. Such theories could posit instead that the ability or inability to be syntactically separated is driven by the two compound classes having distinct kinds of syntactic structure (cf. Zhang, 2007). Regardless of which analysis turns out to be correct, what is important for our first comparison is just that the two types of compounds differ underlyingly in their syntactic representation even when they are not actually separated, and thus that differential neural responses to these two compound types can be attributed to syntactic inference processes.

### 1.2. Compound and simplex nouns in Mandarin Chinese

Although Chinese is sometimes mistakenly characterized as a language “without much morphology”, in fact compound forms dominate the vocabulary of modern Chinese, in some estimates as much as 88% of Chinese lexicon (Zhou & Marslen-Wilson, 1994). Apart from separable vs. inseparable verbs that differ in syntactic structure complexity, in this study we also introduced a second pair of conditions in which the items differ in morphological structure complexity, but do not differ in their ability to be visibly separated in the syntax. This second comparison made use of transliteration-based loanwords in Mandarin like *pi1sa4* (from “pizza”), which are unambiguously simplex (monomorphemic according to lexicalist theories; cf. Hsu et al., 2019; Wei et al., 2023). We compared the ERP (event-related potential) response to these simplex nouns with inseparable compound nouns to yield what would be traditionally considered a “morphological structure” contrast, and we then conducted a topographical similarity analysis to evaluate whether this effect is qualitatively similar to the “syntactic structure” contrast between separable and inseparable verbs, as might be predicted by “syntax-all-the-way-down” theories.

In the current study, we focused on ERP responses in the 275:400ms time-window associated in prior literature with LAN (left anterior negativity) modulations of syntax and morphology. Particularly relevant for our current study, Wei et al. (2023) reported results from a contrast between Mandarin compounds and simplex words (with a variety of part-of-speech, including loanword nouns as in our current study) showing an increased LAN response for compounds relative to simplex words in the 275:400 ms time window. Although we similarly expected to see differential effects of structure over left anterior electrodes in the current study, we chose to be cautious here by testing for effects in multiple regions across the head, because the EEG system used in this study required a different referencing technique (an average reference) that could potentially cause differences in scalp distribution. Complementary to this analysis, we also employed topographical similarity analyses to test whether and when the two complexities modulates scalp distribution in similar ways. Since prior studies suggested that the concreteness of nouns could also modulate anterior channels in a similar time window (where more concrete nouns are more negative than less concrete nouns in amplitude; Zhang et al., 2006; Adorni & Proverbio, 2012; Barber et al., 2013), participants also rated the concreteness of compound and simplex nouns after the experiment. In the current study, we embedded all linguistic expressions in relatively neutral sentential contexts (Table 2), in order to ensure more realistic processing compared to isolated-word paradigms (see Section 3.2).

**Table 2.** Example sentential stimuli for our experiment; the critical segment (i.e., the “critical word”) for our analysis is marked in bold.

|  |
| --- |
| **Compound and simplex nouns** |
| **Structure of sentences** |
| [Because] **[compound/simplex noun]** [is] [segment 4] [segment 5,] [therefore/so] [continuation.]<br>[因为] **[compound/simplex noun]** [是] [segment 4] [segment 5,] [所以/因此] [continuation.] |
| **Example:** |
| 因为/**热狗**/是/一种/高热量的/食物, /所以/不是很健康。<br>Because/**hotdog**/is/a.kind.of/high-calorie/food,/therefore/not.very.healthy.<br>“Because hotdog is a kind of high-calorie food, therefore it is not very healthy.” |
| **Separable and inseparable verbs** |
| **Structure of sentences** |
| [Name] [wants to/doesn't want to/really wants to] **[separable/inseparable verb,]** [so/therefore] [continuation.]<br>[Name] [想要/不想/很想] **[separable/inseparable verb,]** [所以/因此] [continuation.] |
| **Example:** |
| 小范/很想/**造反**, /所以/他秘密组织了一支军队。<br>Fan[name]/really.wants.to/**rebel**,/therefore/he.gathered.an.army.secretly.<br>“Fan really wants to rebel, therefore he gathered an army secretly.” |

## 2. Results

### 2.1. Behavioral results

#### Accuracy for the main experiment

The analyzed subjects generally had a high accuracy in the continuation congruity judgements (mean 96%, range 85%-100%), suggesting that they were attentively comprehending the sentences.

#### Post-test ratings

The separability ratings results confirmed that the separable verbs we used were indeed more separable than the inseparable verbs (separable: 6.1; inseparable: 2.5; *t*(23) = 20.15, *p* < 0.001, two-tailed). For the concreteness ratings, all item types received relatively high concreteness ratings, but we did observe a numerically small but significant difference between conditions (compound: 6.3, simplex: 6.2, *t*(23) = 3.93, *p* < 0.001, two-tailed). For details of these post-tests see the Methods section.

### 2.2. ERP analysis: the LAN time window (275:400 ms)

We conducted an omnibus repeated-measure ANOVA analysis, with ROI (5 ROIs, Figure 2a), contrast type (the morphology contrast, the syntax contrast), and complexity (more complex, less complex morpho/syntactic structure) as independent variables, and the mean response within the 275:400 ms time window in each ROI as the dependent variable (thus, a 5 × 2 × 2 three-way ANOVA). For example, the compound noun condition corresponds to the contrast type of “morphology” and a complexity of “more complex”.

**Figure 2.**
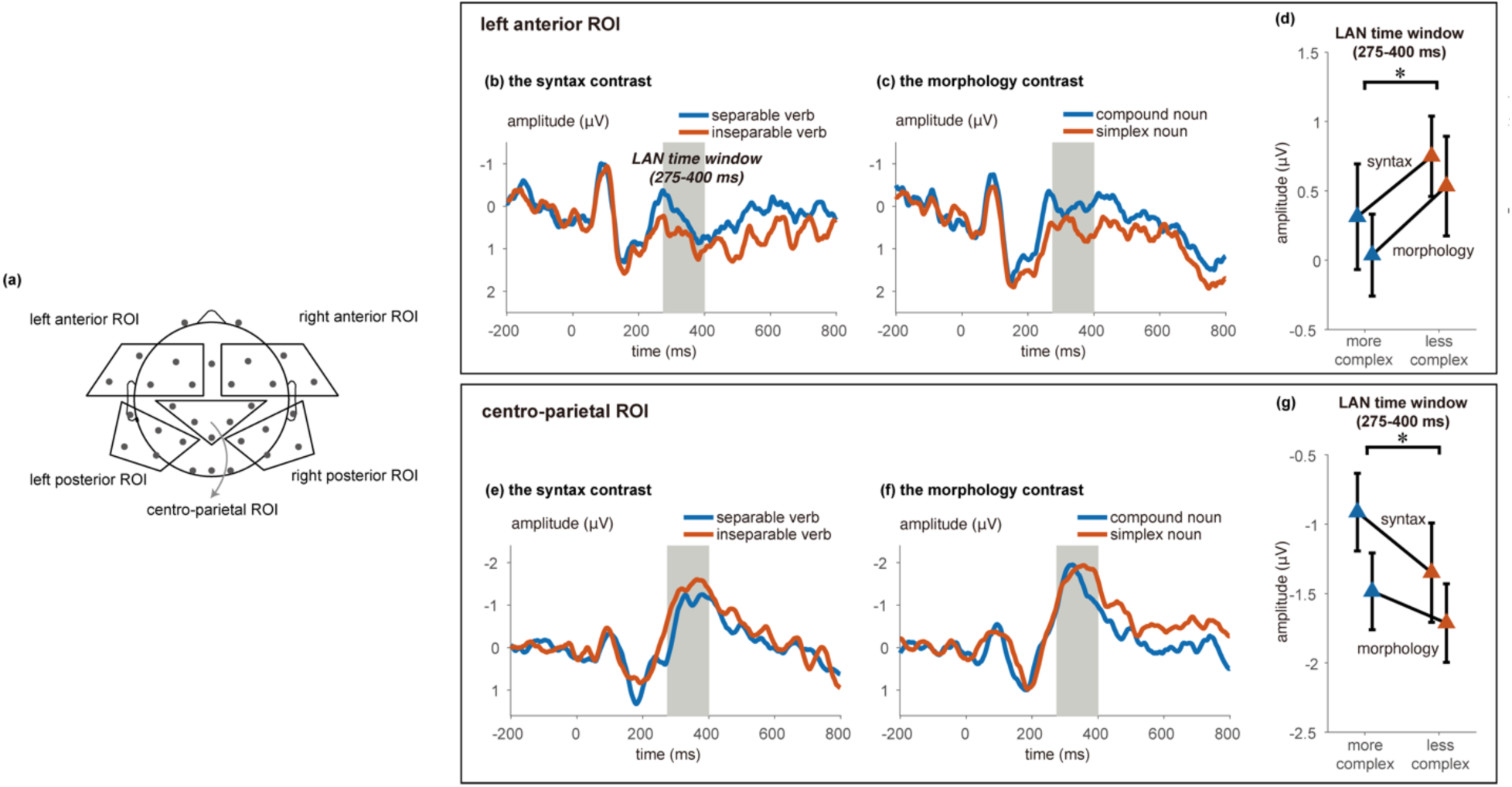
ERP results. ROI definitions are illustrated in (a), and (b) and (c) illustrate the ERPs for all four conditions in the two ROIs which showed significant effects of complexity. **(a)** The ROIs (regions of interest) in our current study. The left anterior ROI consists of F3, F7, FC5, FC7, and FT9; the right anterior ROI consists of F4, F8, FC2, FC6, and FT6; the centro-parietal ROI consists of CP1, CP2, Pz, C3, and C4; the left posterior ROI consists of T7, CP5, P7, P3, and TP9; the right posterior ROI consists of T8, CP6, T8, P4, and TP9. **(b)** Grand average ERPs within the left anterior ROI −200:800 ms around the onset of critical word for compound nouns and simplex nouns. The a priori time window of 275:400 ms is highlighted in gray. **(c)** Grand average ERPs within the left anterior ROI −200:800 ms around the onset of critical word for separable verbs and inseparable verbs. The a priori time window of 275:400 ms is highlighted in gray. **(d)** Mean response and standard error within the 275:400 time window in the left anterior ROI for the “syntax” and “morphology” contrasts (*: *p* < 0.05). **(e)** Grand average ERPs within the centro-parietal ROI for compound nouns and simplex nouns. **(f)** Grand average ERPs within the centro-parietal ROI for separable verbs and inseparable verbs. **(g)** Mean response and standard error within the 275:400 time window in the centro-parietal ROI for the “syntax” and “morphology” contrasts (*: *p* < 0.05). For the ERP waveforms of all scalp electrodes, see Supplementary Materials.

The three-way ANOVA revealed a significant main effect of ROI, *F*(2.07,47.55) = 8.50, *p* < 0.001, a marginally-significant interaction effect between contrast type and ROI, *F*(2.76,63.58) = 2.46, *p* = 0.076, and critically, a significant interaction effect between complexity and ROI, *F*(2.90,66.59) = 2.77, *p* = 0.050; all other effects were not statistically significant (*p’s* > 0.21).

The significant interaction between complexity and ROI suggests that structural complexity indeed modulated neural responses in at least some regions of the scalp. In order to better understand the topography of the structural complexity effect and confirm that it was similar for both morphology and syntax, we followed up the significant interaction with a 2 (contrast type) × 2 (complexity) repeated-measure ANOVA for each of the five ROIs separately. For the left anterior ROI (Figures 2b, d), we indeed observed a significant main effect of complexity, *F*(1,23) = 7.47, *p* = 0.012; the main effect of contrast type and the interaction effect were not significant (*p’s* > 0.19). For the centro-parietal ROI (Figures 2c, e), we also observed a significant main effect of complexity, *F*(1,23) = 6.83, *p* = 0.016, along with a main effect also for contrast type, *F*(1,23) = 6.14, *p* = 0.021, but not an interaction effect, *F*(1,23) = 1.12, *p* = 0.30. For the other three ROIs, no statistically significant effects were observed (*p’s* > 0.14). In all, the ERP analyses seemed to suggest a left anterior effect accompanying a centro-parietal effect in the LAN time window (275:400 ms), which is in line with visual inspection of the temporal evolution of the scalp distributions (Figure 3).

**Figure 3.**
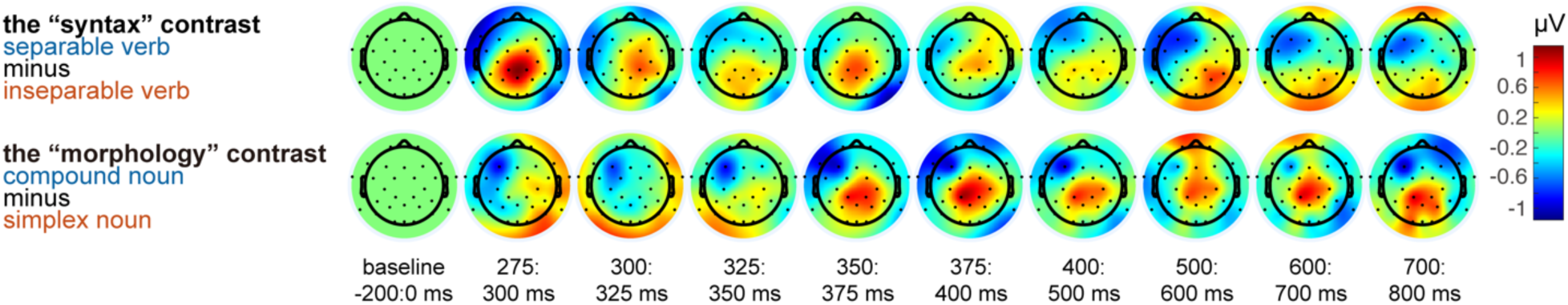
The scalp distribution for the baseline period (−200:0 ms), as well as for every 25 ms between 275:400 ms, and for every 100 ms between 400:800 ms. Here we plot scalp distribution 275:400 ms in smaller (i.e., 25 ms) steps to be more comparable with the results of our topographical similarity analysis.

One remaining question is whether differences in noun concreteness contributed to the effect observed in the morphology contrast for the noun conditions. Although the mean difference in concreteness ratings was quite small (Δ .19 on a 1-7 Likert scale), it was statistically reliable, and therefore it is possible that differences in concreteness rather than morphological status contributed to the difference between compound and simplex nouns. In order to evaluate this possibility, we conducted follow-up tests leveraging individual differences in concreteness ratings (cf. Xu & Li, 2020). If the greater LAN for compound nouns were due to concreteness, we would expect the participants who rated the compound nouns as more concrete than the simplex nouns to demonstrate a more pronounced LAN effect relative to participants who rated the compound nouns as similarly concrete as (or event less concrete than) simplex nouns. This would predict a negative correlation with the LAN effect across participants. However, this correlation was not significant, and numerically went in the opposite direction of what would be predicted (Pearson’s *r* = 0.32, *p* = 0.12). Therefore, we believe it is unlikely that the small difference in concreteness ratings contributed to the ERP differences observed in the compound noun – simplex noun contrast.

### 2.3. Topographical similarity analyses

The second component of our analyses was aimed at explicitly testing the topographical similarity of the “syntax” and “morphology” contrasts across time. This test identified at what points in time the scalp topographies for the two contrasts were similar, in increments of 25 ms (for details see the Methods section). The analysis first identified potential clusters with a *p* < 0.05 threshold, and retained only clusters that survived an α = 0.001 threshold with FDR (false discovery rate) correction. Significant clusters are marked with black contours in Figures 4a, b, for the angle (cosine) similarity analysis and projection amplitude analysis respectively. Four significant clusters were identified in the angle similarity analysis, spanning [50:125 ms (syntax), 450:475 ms (morphology)], [300:375 ms, 425:500 ms], [250:375 ms, 350:400 ms], and [600:700 ms, 175:275 ms] respectively. Three significant clusters were identified in the projection amplitude analysis, spanning [300:375 ms, 450:475 ms], [275:350 ms, 300:400 ms], and [525:625 ms, 225:300 ms] respectively. Critically, both analyses revealed a cluster at 350 ms across contrasts, in line with our LAN time window.

**Figure 4.**
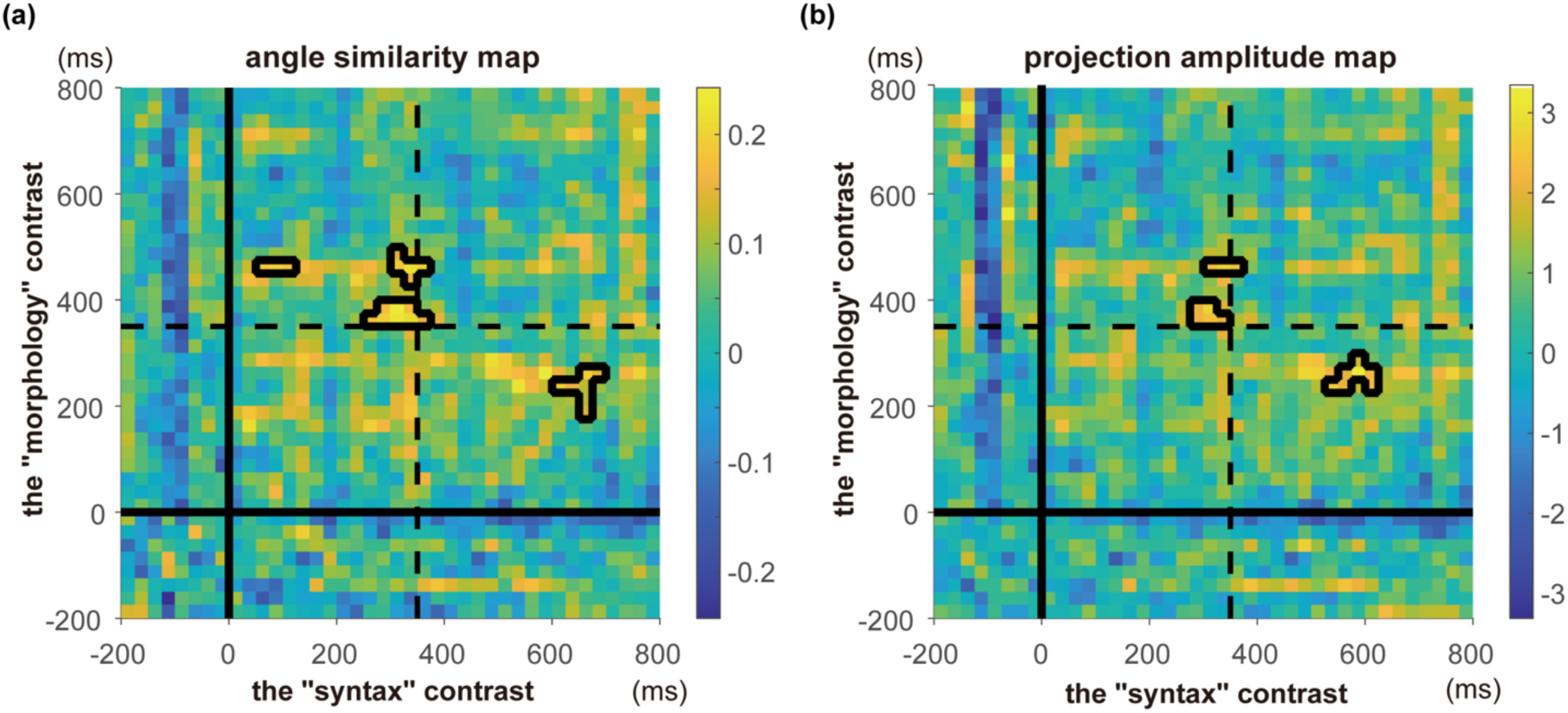
**(a)** Angle (cosine) similarity map from the topographical similarity analysis. The significant (*p* < 0.001, FDR corrected) clusters are outlined in black. The solid lines represent the onset of critical stimuli; the dashed lines indicate 350 ms post-stimulus presentation. **(b)** Projection amplitude map from the topographical similarity analysis. The significant (*p* < 0.001, FDR corrected) clusters are outlined in black. The solid lines represent the onset of critical stimuli; the dashed lines indicate 350 ms post-stimulus presentation.

Interestingly, for the cluster around 350 ms, similar topographies appeared to emerge earlier for the syntax contrast than for the morphology contrast. This is in line with visual inspection of the scalp distributions around 350 ms (Figure 3). Indeed, the exact time course for morphosyntactic structure building has been found to be affected by linguistic factors (Slaats et al., 2024).

## 3. Discussion

In the current study, we observed that both syntactic complexity and morphological complexity modulated left anterior and centro-parietal electrodes in the same direction at 275:400 ms. The similarity of the neural response to the two contrasts was corroborated by topographical similarity analyses which identified a cluster of similarity around ∼350ms. The similarity in the ERP responses for the morphological and syntactic contrasts is in line with the hypothesis that there is no qualitative difference between morphological and syntactic structure building (the “syntax-all-the-way-down” hypothesis; Oseki & Marantz, 2020; Krauska & Lau, 2023).

### 3.1. Neural responses to separable verb compounds

In the current study, we observed left anterior and centro-parietal effects for the “syntax” contrast, i.e., between separable and inseparable verbs, in the 275:400 ms time window. To restate the linguistic fact, the separable verbs differ from inseparable verbs in that they are separable by other syntactic units, suggesting that separable verbs correspond to a more complex syntactic structure compared to inseparable verbs (see Section 1.1). Because both types of verbs are semantically non-compositional and consist of two morphemes, these neural effects should be attributed to the inference of *syntactic* structure.

Several previous EEG studies have also examined Mandarin separable verbs (Zhang & Jiang, 2010; Gu, Yu, et al., 2011; Gu, Yang, et al., 2018), but with a different goal and a correspondingly different approach. As these studies assumed a lexicalist architecture, they asked whether separable verbs are “words” or “phrases” by comparing whether the ERP to separable verbs was more similar to words or to phrases. Zhang and Jiang (2010) found that separable verbs differed from words in many components (P200, N400 and P600) but from phrases only in P600; however, Gu et al. (2018) failed to observe a significant effect for the P600. These mixed results may have arisen due to additional factors in the materials these studies used for comparison; it is unclear whether all the items in the “word” conditions were verbs, and it is also unclear whether transitivity was matched across conditions. Furthermore, although aimed at distinguishing differences between “word” and “phrase” processing, these studies employed lexical decision or semantic categorization tasks, in which phrase comprehension is less likely to proceed normally. To our knowledge no prior study has compared separable and inseparable verbs within sentence contexts while matching their transitivity, which we take to be crucial for evaluating their processing differences.

Since we used an average re-reference rather than the mastoid re-reference used by most language EEG studies, we need to be cautious about making assumptions about the relationship between the topography of our ERP effects and those observed in prior studies. Broadly speaking however, our current left anterior and centro-parietal effects in the LAN time window are in line with a previous ERP study which observed a LAN effect in a manipulation of Dutch particle verbs (Piai et al., 2013; with averaged mastoid re-reference). Dutch particle verbs consist of a particle and a head verb which together can carry an idiomatic meaning, and as in the case of Mandarin separable verbs, they preserve this meaning even when they are syntactically separated^3^. For example, the particle verb *voor-spellen* (before-mirror, “predict/promise”) can be separated, with the particle preceding the head verb:

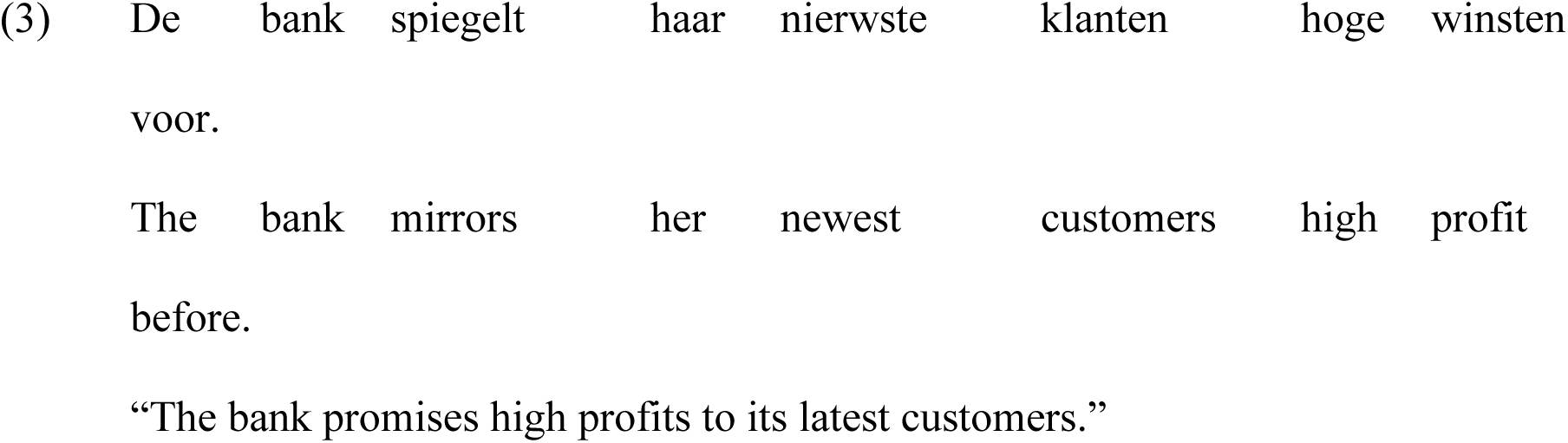

Piai and colleagues compared the ERP responses for the head verbs that could appear in particle verbs against “inseparable” verbs that could not form a particle verb with any particle, when they were embedded in sentences. They hypothesized that, although the head verbs could also stand alone, participants would still consider the more complex particle-verb syntactic structure already at the head verb (particularly since participants encountered many particle verbs in the experiment). Indeed, they observed a LAN effect in which the head verbs which could participate in particle verb constructions exhibited a more negative effect in left anterior electrodes between around 300-550ms relative to the verbs that could not, consistent with the hypothesis that the more complex syntax of the particle verb construction put greater demands on working memory. One alternative explanation for this effect was that it could simply reflect greater syntactic uncertainty, although Piai et al. (2013) considered this explanation less likely since they failed to see a difference in the LAN response between head verbs that could take a wider range of particles vs. a smaller range. The current study convergingly suggests that mere syntactic structure inference/building is enough to elicit a LAN-like effect, as we presented both parts of intransitive separable verbs at once to the subjects, minimizing the role of syntactic prediction.

Although both Mandarin separable verbs and Dutch/German particle verbs are separable, there appears to be intriguing differences between the two phenomena, which would serve as a fertile ground for future research. Interestingly, for Mandarin separable verbs, the type of syntactic components that could go in between, as well as the syntactic operations that they could undergo (e.g., movement, reduplication) seem to be quite idiosyncratic. As an example, consider 帮忙(bang1mang2, [help]-[busy], “help”) and 费心 (fei4xin1, [cost]-[heart], “take trouble”). Although they are both separable verbs, they exhibit different acceptability profiles when separated by certain syntactic units, and when undergoing movement. While bang1mang2 can be separated in 帮了一个大忙 (bang1le0yi2ge0da4mang2, `bang1` ASP one CL big `mang2`, “did a huge favor”), 费了一个大心 (fei4le0yi2ge0da4xin1, `fei4` ASP one CL big `xin1`, intended “took huge trouble”) is unacceptable. While 忙他帮了 (mang2ta1bang1le0, `mang2` he `bang1` ASP, “he helped”) is fine, 心他费了 (xin1ta1fei4le0, `xin1` he `fei4` ASP, intended: “he took some trouble”) is tricky. This is just one pair of examples among many Mandarin separable verbs, many syntactic components that could go in between, and many syntactic operations that they could undergo (e.g., movement, reduplication of the first part; see Siewierska et al., 2010; Wang, 2011). This interesting idiosyncrasy remains to be documented comprehensively. How do native speakers come to learn this idiosyncrasy? Are there common factors that could account for the idiosyncrasies in Mandarin separable verbs and in German/Dutch particle verbs (see Trotzke et al., 2015; Schoenmakers & Foolen, 2022)? All these interesting questions await future research.

A separate question is where in the brain the ERP effects originates from, and what that indicates about its functional interpretation. Many MEG studies have also tried to single syntax out (i.e., controlling for factors other than syntactic structure complexity) in order to examine brain responses sensitive to syntactic structures, and they have observed effects in the posterior temporal lobe in somewhat similar time windows (Flick & Pylkkänen, 2020; Law & Pylkkänen, 2021; Matar et al., 2021). For example, Law and Pylkkänen (2021) examined MEG responses in the left posterior temporal lobe (PTL) when a list was embedded in a sentence vs. a longer list, and observed a difference in the 333-389 ms time window. This is also in line with many fMRI studies supporting the involvement of the posterior temporal lobe in syntactic structure inference (Matchin et al., 2017; Zaccarella et al., 2017; Matchin & Hickok, 2020). We thus speculate that our ERP effects may originate from the posterior temporal lobe, which remains to be confirmed by future studies with techniques with a better spatial resolution (e.g., MEG and fMRI).

Although we have argued that our current ERP effect reflects sensitivity to syntactic structure, we should note that it is unclear if the effect reflects the recruitment of stored structured syntactic objects or “active” online structure building. The difference between these two processes is subtle but important. During syntactic structure building in language comprehension, one may consult structured objects (e.g., specific syntactic treelets) stored in long-term memory, but ad hoc syntactic structure building is more than just “reinstantiating” all relevant stored objects. However, it is difficult to empirically disentangle the two processes as of now, because the neural implementation of syntactic structure building remains highly debated (Ding et al., 2016; Ding, 2023; Meyer, 2018; Pylkkänen, 2019; Matchin & Hickok, 2020; Martin, 2020; Murphy, 2020, 2024; Kazanina & Tavano, 2023; Krauska & Lau, 2023), as well as the question of how long-term linguistic objects are stored (Poeppel & Idsardi, 2022; Krauska & Lau, 2023; Murphy, 2024).

### 3.2. Neural responses to Mandarin noun compounds vs. simplex forms

Our left anterior and centro-parietal effects for the “morphology” contrast (i.e., compound vs. simplex forms) in a sentence context can be seen as a replication of Wei et al.’s (2023) earlier findings, with a similar time window. Wei and colleagues employed a lexical decision task, and observed a LAN effect in comparing Mandarin compounds and simplex forms with a variety of parts-of-speech. Our current study extends and strengthens their findings, as the results of isolated-word paradigms do not always agree with the processing dynamics observed when the same items are embedded in sentential contexts (see Morris, 2017). Mirroring this point, a recent MEG study on auditory word processing revealed different neural substrates when words are processed in isolation vs. in continuous speech (Gaston et al., 2022). Concerns have also been raised about the lack of naturalness of isolated-word comprehension paradigms (Salmelin et al., 1999; Hamilton & Huth, 2020). Our corroborating evidence from a sentence paradigm is thus useful in demonstrating that the differential processing Wei and colleagues observed for compound Mandarin forms in lexical decision generalizes to how they are processed in integrated sentence contexts.

A component that has been argued to be sensitive to morphological structure in the MEG literature is the M350 (Stockall & Marantz, 2006; Fiorentino & Poeppel, 2007; Brooks & Cid de Garcia, 2015; Cavalli et al., 2016; Hsu et al., 2019; for reviews see Leminen et al., 2019; Royle & Steinhauer, 2023). The term “M350” sometimes refers to different measurements around 350 ms after the word onset, including the RMS (root mean square) peak around 350 ms (Fiorentino & Poeppel, 2007), as well as neural responses in posterior temporal regions around 350 ms (Brooks & Cid de Garcia, 2015; Cavalli et al., 2016; Hsu et al., 2019). The left anterior and centro-parietal effects observed in our current EEG study may reflect the same neural processes as M350, based on its time window. However, the neural source of the current effects remains to be confirmed by future MEG studies with better spatiotemporal resolution.

### 3.3. Similarity between “syntactic” and “morphological” effects

In short, in the current EEG study we observed a similar LAN effect for both the “syntax” and the “morphology” contrasts. Within the a priori 275:400 ms time window, we observed similar left anterior and centro-parietal effects that are both sensitive to syntactic structure complexity, but also morphological structure complexity, within the same group of subjects. This observation was corroborated by our topographical similarity analyses across the two contrasts, revealing a significant cluster around 350 ms, suggesting that the scalp distribution around 350 ms was similar across the two contrasts. The left anterior and centro-parietal effects may originate from the same neural source, resulting in two opposing adjacent maxima on the scalp.

Form-level visual parsing is considered to happen around 170 ms or earlier (Tarkiainen et al., 1999; Salmelin, 2007, 2010; Gwilliams et al., 2016; Sigurdardottir et al., 2021). Based on the later time window of our effects (275:400 ms), it is thus likely to reflect abstract (i.e., non-visual) morpho-syntactic processes. In fact, even the ∼ 170 ms effects are already affected by morphosyntactic rules (Gwilliams & Marantz, 2018). From the traditional lexicalist point of view, the shared neural response across the “morphology” and “syntax” contrasts may come as a surprise; since a clear cut is assumed between morphological and syntactic rules and representations, the lexicalist view does not predict that they should share linguistic parsing processes, any more than phonology and syntax would be predicted to share linguistic parsing processes. On the other hand, shared process(es) are predicted by non-lexicalist theories of language comprehension (see Oseki & Marantz, 2020; Krauska & Lau, 2023 for review) which hold that there is no qualitative difference between so-called “syntactic” and “morphological” structure inference/building. In other words, syntax is “all the way down”.

Interestingly, in prior EEG and MEG work effects in this time-window (LAN and M350) have been argued to reflect structure inference in the neuro-morphology and the neuro-syntax literature respectively, and both have sometimes been suspected to originate from posterior temporal regions as reviewed above (note that although some studies suggest a left frontal source for LAN effects, many of those cases involved grammatical violations; e.g., Friederici & Kotz, 2003; Carreiras et al., 2010). Our current study verifies that a similar effect for “morphology” and “syntax” can be observed within the *same* group of participants. Of course, a similar topography does not guarantee the exact same neural processes, yet it provides encouraging *prima facie* evidence awaiting future studies with better spatial resolution to confirm.

Indeed, either “morphology” or “syntax” may consist of a variety of different operations that may correspond to non-identical neural signatures. Nevertheless, some neural signatures could be shared across different combinatorics, which may be governed by e.g., the topology of the morpho-syntactic trees (for example those relevant to the depth or node count of the trees, see Brennan et al., 2016; Li et al., 2016; Ohta & Sakai, 2017). These shared neural signatures (suggesting shared neural processes) may be the reason why we observed a similar neural response across the “morphology” and “syntax” contrasts in our current study, regardless of the idiosyncrasy in morpho-syntactic combinatorics.

### 3.4. The opacity conundrum

One question is the degree to which effects like the current ones are modulated by semantic opacity: how much the meaning of the whole is intuitively related to the meaning of the parts. Firstly, we would like to point out that Wei et al. (2023) did not observe a significant modulation effect of opacity on LAN or N400 with Chinese compound nouns of different opacity. Although semantic opacity may still be an interesting factor to explore in the future, we would also like to point out some difficult theoretical challenges regarding “semantic opacity”, which we term “the opacity conundrum”.

To begin with, it is worth noting that there is no independent measurement of *semantic* opacity apart from subjective ratings. For one thing, it is hard, if not impossible, to prove that the measured ratings are purely of semantic nature but not of morpho-syntactic nature at all (**the semantic purity problem**). For another, there is no obvious way to tell if the measured opacity is just a reflection or natural result of different underlying morpho-syntactic structures, or that opacity affects morpho-syntactic parsing in turn (**the direction of causality problem**). Another challenge for the opacity for complex nouns is that, opacity has to be defined somehow by the relations between the meaning of the whole linguistic expression and *the meaning of a morpheme*, which is an extremely ill-defined concept in linguistics (**the morpheme meaning problem**). Notably, expert linguists’ judgments do not match perfectly with novice ratings (Gagné et al., 2016); in this case it is hard to argue for one judgment over the other precisely because of a lack of independent measurement and easily-operationalizable definition for semantic opacity (same for the construct “semantic transparency”). Therefore, although we acknowledge that the interplay between “opacity” and morpho-syntactic parsing is an interesting research question, there are some deep theoretical problems about “opacity” which need to be addressed in future research, namely, the semantic purity problem, the direction of causality problem, and the morpheme meaning problem.

## 4. Conclusion

In the current study, we observed a similar modulation of syntactic and morphological structure complexity on left anterior and centro-parietal electrodes in a time window informed by prior research (275:400 ms). These qualitatively similar responses are in line with the predictions of non-lexicalist theories of language comprehension, which proposes that there is no qualitative difference across the neuro-cognitive operations/computations involved in syntactic and “morphological” structure inference and building.

## 5. Methods

### 5.1. Subjects

Subjects were recruited and tested in Shanghai, China, and all of them self-identified as native speakers of Mandarin Chinese. Subjects received monetary reimbursement for their participation. Written consent was acquired from each subject, and the procedures were approved by the Institutional Review Board of the University of Maryland and New York University Shanghai. Two subjects’ data were excluded from data analysis because of artifacts (e.g., blinking, drifts) in excessive (≥ 50%) epochs. The remaining 24 subjects (16 female), had an age of 19-29 (*M* = 23). One of the analyzed subjects was left-handed, and all others were right-handed, as measured by a translated version of the Edinburgh Handedness Inventory (Oldfield, 1971).

### 5.2. Stimuli

We first selected 30 items for each of the four compound types tested in this study: compound nouns, simplex nouns, separable verbs, and inseparable verbs. The compound and simplex nouns were disyllabic, matched in log10 bigram frequency in two large corpora (BCC multi-domain corpus, Xun et al., 2016; simplex: 3.6, compound: 3.7, independent sample t test *p* = 0.33, two-tailed; CCL2024 corpus, Zhan et al., 2019; simplex: 3.9, compound: 3.9, independent sample t test *p* = 0.88, two-tailed), as well as the stroke numbers for each character (1st character, simplex: 8.1, compound: 8.3, Mann-Whitney U test *p* = 1.0, two-tailed; 2nd character, simplex: 6.9, compound: 7.7, independent sample t test *p* = 0.29, two-tailed; the Mann-Whitney U test was administered instead of the independent sample t test whenever at least one compared distribution did not pass the Shapiro-Wilk test, *p* < 0.05). The simplex nouns were transliteration-based loan words. Although a number of these loan words in Chinese are composed of two visually similar characters, for example, 咖啡 (ka1fei1, “coffee”; the two characters share the same radical to the left), 芭蕾 (ba1lei2, “ballet”; the two characters share the same radical on top), we were careful to exclude such items from our stimuli set. The separable and inseparable verbs were intransitive, disyllabic, and matched in log10 bigram frequency in two large corpora (BCC multi-domain corpus; separable: 3.9, inseparable: 3.9, independent sample t test *p* = 0.92, two-tailed; CCL2024 corpus; separable: 4.2, inseparable: 4.1, Mann-Whitney U test *p* = 0.23, two-tailed), as well as the stroke numbers for each character (1st character, separable: 8.6, inseparable: 8.2, independent sample t test *p* = 0.55, two-tailed; 2nd character, separable: 7.9, inseparable: 8.6, independent sample t test *p* = 0.36, two-tailed). All compounds used in the study were selected to be intuitively non-compositional in meaning, in that the meaning did not follow straightforwardly from the meaning of the parts, e.g. for the compound 热狗 (re4gou3, hot-dog, “hotdog”), its meaning isn’t a straightforward composition of the meaning of 热 (re4, “hot”) and 狗 (gou3, “dog”).

We then created a list of 120 sentences that contained the items from the four 30-word compound sets. The idea was to embed the compound words in a low-prediction neutral context. The sentential patterns and example sentences are illustrated in Table 2. Specifically, for the 30 separable and inseparable verb sentences respectively, 1/3 followed “wants to”, “doesn’t want to” and “really wants to” respectively. In order to control for the semantic congruity of “wants to/doesn’t want to/really wants to + verb” (which all consist of 4 characters) across the two conditions, we matched the 4-gram frequency in two large corpora (cf. Lau et al., 2016). Because many of the 4-grams did not appear in the corpus (despite them being large of their kind), we calculated log10(frequency+1) as log10 frequency and employed a Mann-Whitney U test; the two conditions were balanced in 4-gram frequency (BCC multi-domain corpus: separable: 0.39, inseparable: 0.34, *p* = 0.58; CCL2024 corpus: separable: 0.38, inseparable: 0.43, *p* = 0.77). Two balanced lists of names were created for the separable and inseparable verb sentences and the names were counter-balanced across the 24 analyzed subjects. For separable and inseparable verb sentences, a comma always followed the verbs in order to ensure an intransitive reading, because of an increasing trend in Mandarin where intransitive verbs can be somewhat acceptable with transitive usages (Liao & Tsai, 2023). An additional 60 filler sentences were adapted from another study (Liao & Lau, 2020), rendering 180 sentences for each subject.

### 5.3. Procedure

Subjects sat ∼ 60 cm in front of the screen. 180 trials were run for each subject (30 trials for each of the 4 conditions + 60 filler sentences), with a different sentence presented in each trial. Each trial began with a fixation cross of 1000 ms, followed by presentation of the segments of this sentence in a segment-by-segment (i.e., RSVP, rapid serial visual presentation) manner, with a presentation duration for 600 ms for each segment, and a 200 ms interval between segments, for a total 800ms stimulus-onset asynchrony (Figure 1b). The segments were presented in white characters against a black background, in 60 pt, Yahei font; they were presented at the center of the screen. In 1/3 of the trials, the last segment of the sentence (i.e., the continuation) was colored in green and the subjects were instructed to respond to whether this continuation was congruent in this sentence, by pressing m (incongruent) and n (congruent) on the keyboard with their right hand (the responses did not have a time-limit). In the other 2/3 of the trials, the last segment was presented on the screen in white briefly for 1000 ms, and subjects did not need to respond. The intertrial intervals were self-paced; subjects pressed the space bar when they were ready to start the next trial.

After the experiment, the subjects were administered two post-tests. The first, “separability” post-test aimed at testing their subjective rating of the separability of the 60 separable and inseparable verbs used in the experiment. Subjects rated the acceptability of the separable and inseparable verbs on a 1-7 Likert scale, when they were separated by aspect markers “了” (le0) or “过” (guo4). The aspect marker used for each verb in the post-test was balanced across subjects; for example, half of the subjects were asked to rate the acceptability of “造了反” ([rebel-] le0 [-rebel]), and the other half of the subjects were asked to rate the acceptability of “造过反” ([rebel-] guo4 [-rebel]). Separability by aspect markers is a common and signature property for separable verbs (Siewierska, Xu & Xiao, 2010; Wang, 2011). The second post-test asked subjects to rate the concreteness of the 60 compound and simplex nouns used in the experiment on a 1-7 Likert scale, with an instruction adapted from prior concreteness studies in Chinese (Yao et al., 2017; Xu & Li, 2020).

### 5.4. Behavioral data analysis

#### Accuracy for the main experiment

Subjects’ responses to the continuation judgement task were recorded, and we calculated the accuracy for each subject.

#### Post-test ratings

For the separability and concreteness ratings, we conducted pairwise t-tests on the mean ratings for each subject by condition.

### 5.5. EEG recording

The EEG was recorded using a 32-channel (Fp1, Fz, F3, F7, FT9, FC5, FC1, C3, T7, TP9, CP5, CP1, Pz, P3, P7, O1, Oz, O2, P4, P8, TP10, CP6, CP2, C4, T8, FT10, FC6, FC2, F4, F8, Fp2, Cz) active electrode system (Brain Vision actiCHamp; Brain Products) with a 1000 Hz sampling rate in an electromagnetically shielded and sound-proof room. Electrodes were placed on an EasyCap, on which electrode holders were arranged according to the 10-20 international electrode system. The impedance of each electrode was kept below 25 kΩ. The data were referenced online to electrode Cz. Two additional EOG electrodes (HEOG and VEOG) were attached for monitoring ocular activity. The EEG data were acquired with Brain Vision PyCoder software and filtered online between DC and 200 Hz with a notch filter at 50 Hz.

### 5.6. EEG data analysis

EEG data were analyzed with EEGLAB and ERPLAB, along with customized MATLAB scripts. The EEG data was first subject to a 0.01 Hz to 40 Hz band-pass filter. Then bad channels were interpolated (spherical, 0-2 bad channels per subject), and the data was re-referenced to the average reference across scalp electrodes. An average re-reference was used because we did not have mastoid channels in our EEG setup. Then, −200:800 ms epochs were extracted and baselined to the mean response between −200:0 ms. Epochs with artifacts were identified and excluded with a simple voltage threshold of −100 to 100 μV. We then reviewed the data, and if needed, adjusted the voltage threshold for individual subjects. A few trials were excluded due to a technical error that resulted in an inaccurate presentation duration. Then the ERPs for each condition were calculated within each subject. The number of trials left for each condition after artifact rejection is as follows: simplex nouns (26.3 trials, range 18-30), compound nouns (26.7 trials, range 21-30), separable verbs (27.3 trials, range 19-30), inseparable verbs (27.0 trials, range 19-30).

#### ERP (event-related potential) analysis

We conducted an omnibus repeated-measure ANOVA analysis, with ROI (5 ROIs, Figure 1c), contrast type (the morphology contrast, the syntax contrast), and complexity (more complex, less complex morpho/syntactic structure) as independent variables, and the mean response within the 275:400 ms (Wei et al., 2023) time window for each subject in each ROI as the dependent variable (thus, a 5 × 2 × 2 three-way ANOVA). Greenhouse-Geisser correction was applied whenever sphericity was violated. The ANOVA analyses were conducted in JASP 0.18.1 (JASP Team, 2023).

#### Topographical similarity analysis

Topographical similarity analysis (i.e., representational similarity analysis, spatial similarity analysis) is an analysis for comparing the topographical distribution of ERP responses that is free from bias in electrode selection (Tian & Huber, 2008; Murray et al., 2008; Tian et al., 2010; Wang et al., 2019). Quantifying topographical similarity with this method has been used to hallmark similarity in the underlying neural processes in many language comprehension studies (Yang et al., 2020; Wang et al., 2020; Wang & Kuperberg, 2023; Hubbard & Federmeier, 2021; Wei et al., 2023; Zhao et al., 2023; Huang et al., 2023; Ding et al., 2024).

For a more stable topography, we first averaged the ERP responses every 25 ms for each subject within the −200:800 ms epochs, collapsing 1000 time points (−200:800 ms) to 50 time points. Then we obtained the scalp topography for the “morphology” contrast (compound minus simplex nouns) and the “syntax” contrast (separable minus inseparable verbs) for each subject respectively. Then, we calculated the angle (cosine) similarity and projection amplitude (Tian & Huber, 2008; Tian et al., 2010) across the two contrasts at each of the 20 ms time windows. We treated the ERP amplitudes of each scalp electrode (31 in total) as an unique dimension of a vector. Suppose the ERP amplitudes for the “morphology” contrast at time window i forms an 1×31 vector **A_i_**, and the ERP amplitudes for the “syntax” contrast at time window j forms an 1×31 vector **B_j_**, the angle similarity of the two vectors was calculated as:

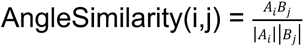

And the projection amplitude of the two vectors was calculated as the mean of the projection of vector **A_i_** on **B_j_** and the projection of vector **B_j_** on **A_i_**:

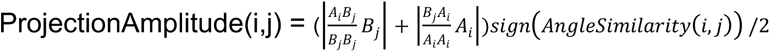

We calculated the angle similarity and projection amplitude for every time window of the “morphology” contrast and each time window of the “syntax” contrast. Thus we obtained a 50×50 map for angle similarity and projection amplitude for each subject respectively. First, at each time window (i,j) in the angle similarity map (which contains one single angle similarity value for each subject), we conducted a one-sample t test against 0 (that the two topographies are orthogonal to each other, i.e., not similar at all), *p* < 0.05 (uncorrected). Then a set of potential clusters were identified based on temporal adjacency of these significant time windows. After that, for each potential cluster, we calculated the mean within it for each subject, and compared this distribution against 0 (i.e., no similarity) with a one-sample t test, rendering a new *p* value for each temporal cluster. Only the clusters that survived an α = 0.001 threshold (FDR corrected with the fdr_bh() function, Benjamini & Hochberg, 1995; Groppe, 2023) are outlined in Figure 1c. A similar analysis was run for the projection amplitude map as well (Figure 1d).

## Acknowledgements

We would like to thank Alexander Williams, Clara Cuonzo, David Embick, Xiaoyu Yang, Yanan Du, Jiaqiu Sun and Cas Coopmans for discussions and comments; Yanan Du, Binyan Hu and Xiaoyu Yang for comments on experimental materials; Yuhan Lu, Yuchunzi Wu, Jiaqiu Sun, Hao Zhu and other Tian Lab members for assistance in data collection.

## Author contributions

**Xinchi Yu**: Conceptualization, data curation, investigation, formal analysis, methodology, software, visualization, writing-original draft, writing-review & editing. **Sebastián Mancha**: Writing-review & editing. **Xing Tian**: Resources, supervision, writing-review & editing. **Ellen Lau**: Conceptualization, funding acquisition, methodology, supervision, writing-original draft, writing-review & editing

## Footnotes

1 Note that the analogy is inexact as English doesn’t have verb-verb compounds; this is only meant to give an intuition.

2 By morphological or syntactic “representations”, we refer to the mental representations for the morpho-syntactic nodes and the relations between/among each other in morpho-syntactic trees.

3 Idiomatic phrasal verbs in English (e.g., “heat up”; Cappelle et al., 2010; Paulmann et al., 2015) might be somewhat similar as well.

