## Supplementary Materials for "Shared neural computations for syntactic and morphological structures: evidence from Mandarin Chinese"

**Supplementary Materials: Supplementary Figures**

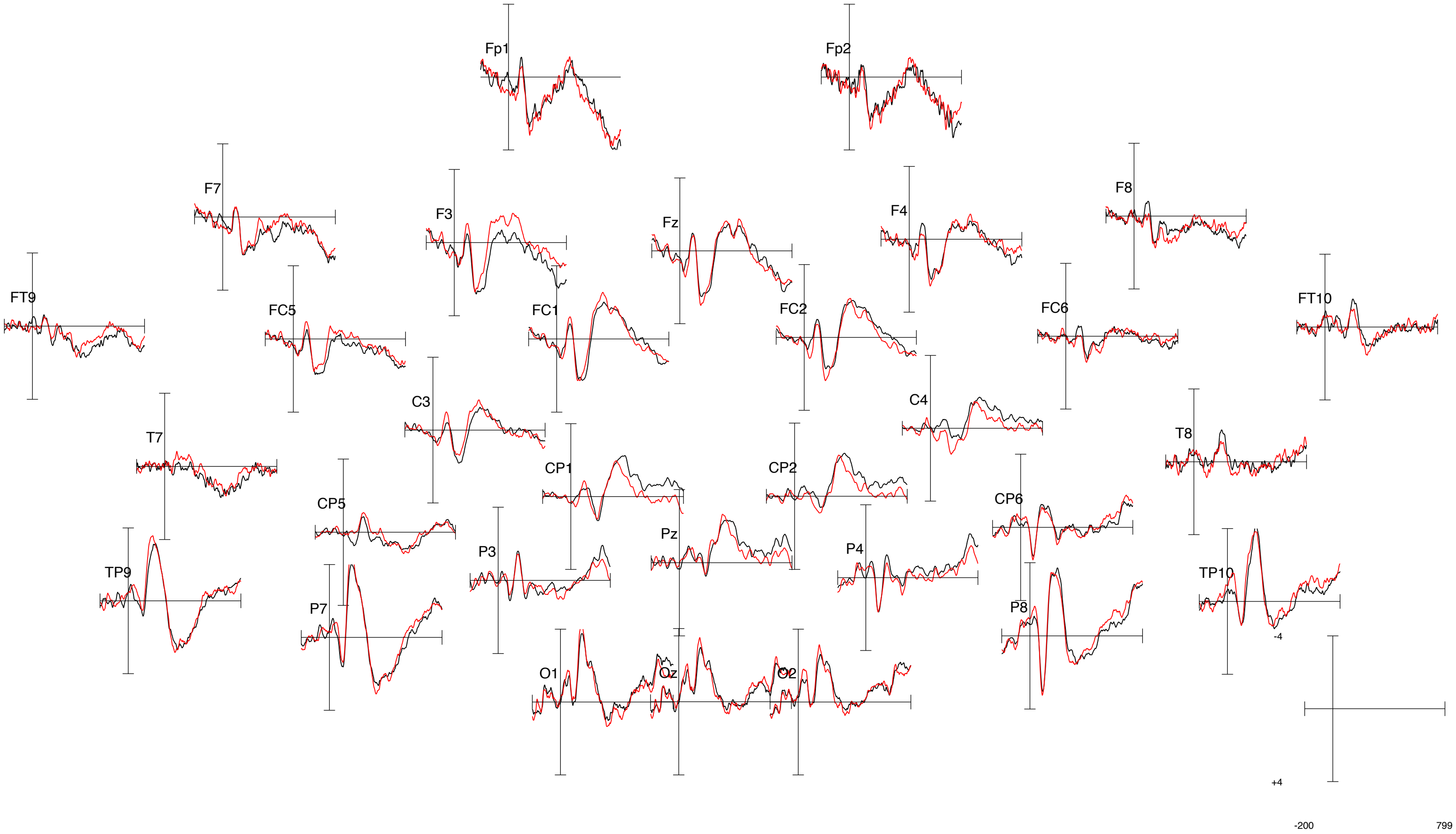


**Figure S1.** ERP waveforms of all scalp electrodes for the "morphology" contrast. Red: compound nouns; black: simplex nouns.

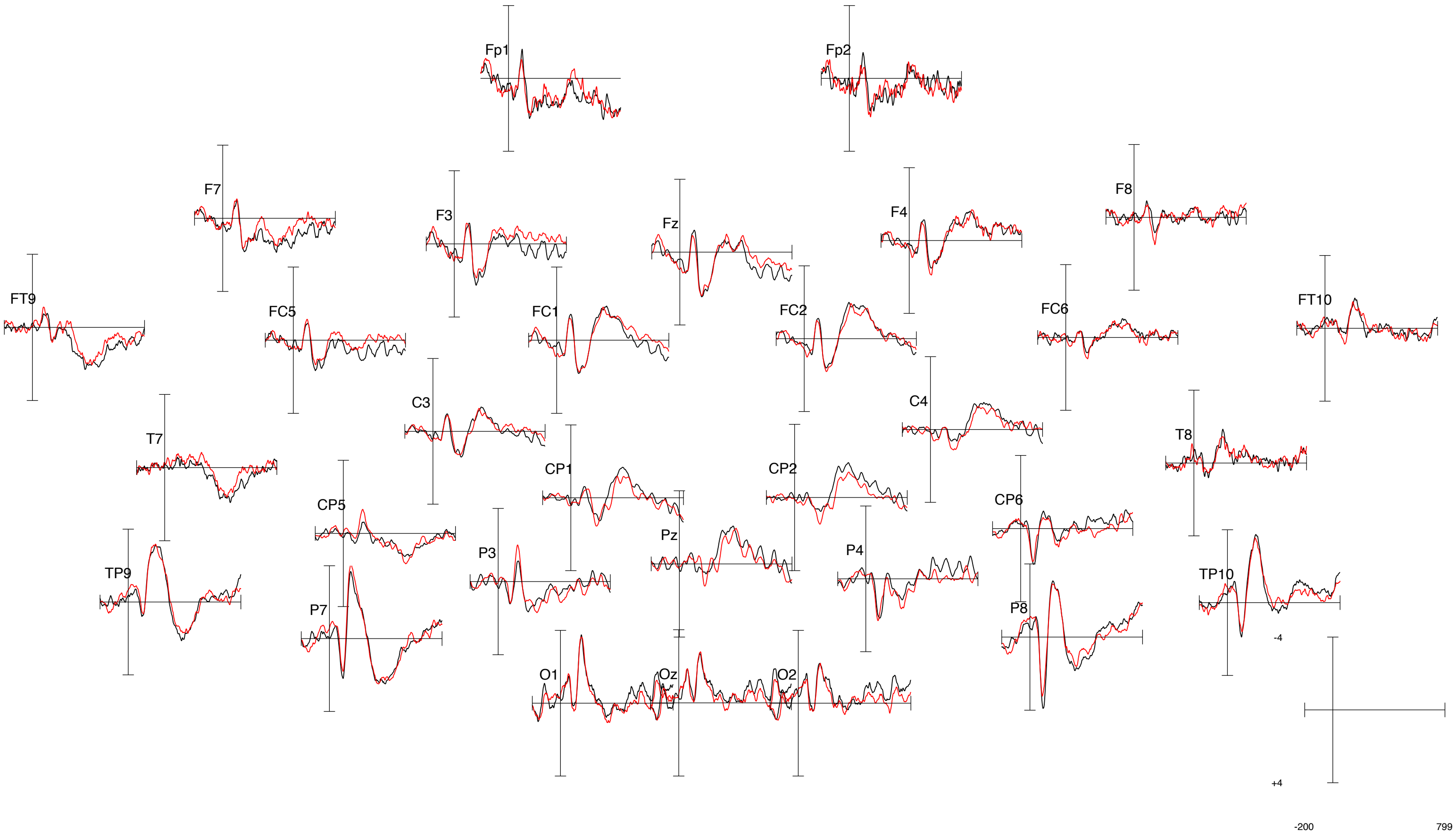


**Figure S2.** ERP waveforms of all scalp electrodes for the “syntax” contrast. Red: separable verbs; black: inseparable verbs.
